# *In Vitro* Efficacy of Paclitaxel-loaded PLGA Nanoformulations for Lung Cancer Treatment Demonstrated by Label-free Multiphoton-Fluorescence Lifetime Imaging Microscopy

**DOI:** 10.64898/2026.06.01.729214

**Authors:** Lia Santos, Maria L. Ribeiro, Tiago Magalhães, Jana B. Nieder, Christian Maibohm

**Affiliations:** INL—International Iberian Nanotechnology Laboratory, Nieder Group on Ultrafast Bio- and Nanophotonics, Av. Mestre José Veiga s/n, 4715-330 Braga, Portugal; Instituto de Investigação e Inovação em Saúde (i3S), Universidade do Porto, Rua Alfredo Allen 208, 4200-135 Porto, Portugal; Instituto Universitário de Ciências da Saúde (IUCS-CESPU), Rua Central de Gandra 1317, 4585-116 Gandra, Portugal

**Keywords:** PLGA nanoparticles, Label-free metabolic imaging, Drug delivery, Cancer therapy, Nanomedicine

## Abstract

Lung cancer remains the leading cause of cancer-related mortality, and despite its limited efficacy, and low specificity, chemotherapy is still commonly used as a first-line treatment. Herein, we prepare and characterize poly(lactic-co-glycolic acid) (PLGA) nanoparticles (NPs) for the use as a non-targeted nanocarrier drug delivery system for the chemotherapeutic drug, paclitaxel (PTX). Drug loading capability, mono-dispersity, zeta potential, morphology, and drug release profile were determined for the PLGA NPs. Furthermore, the IC_50_ of the loaded nanoparticles was determined to be 20 times lower than the IC_50_ value of free PTX in cytotoxic studies in A549 lung cancer cells. The time dependent therapeutic efficacy of both free and encapsulated PTX was assess at several time points in A549 monolayers via label-free multiphoton metabolic imaging based on the fluorescence lifetime of the metabolic cofactor NAD(P)H. Specifically, the significant continuous metabolic shift towards oxidative phosphorylation in cells treated with NPs was correlated with the NPs drug release profile, highlighting the sustained and controlled behavior of the drug delivery system. These findings underscore not only the utility of PLGA NPs as a drug delivery system with enhanced efficacy compared to free drug treatments but also the use of label-free multiphoton fluorescence lifetime imaging microscopy for non-invasive imaging, allowing the tracking of metabolic changes on cellular and subcellular level. Additionally, the presented method is extended to include more complex *in vitro* models allowing for assessing the cellular bioenergetic response of individual cells in 3D models in real time, paving the way for improved therapeutic strategies against lung cancer.

## Introduction

Currently, lung cancer has the highest incidence and mortality worldwide, accounting for a significant proportion of cancer-related deaths. ^1^ Chemotherapy continues to be the first line of treatment for non-small cell lung cancer (NSCLC), used for both early and metastatic cancer states. ^2^ Paclitaxel (PTX) is a well-established drug that is widely used for NSCLC treatment. However, it’s limited efficacy, low specificity, and acquired cellular resistance, underscores the need for innovative drug delivery systems that can enhance the therapeutic outcome. ^3^ Nanoparticle-based drug delivery systems can overcome these limitations by improving drug solubility, stability, and release kinetics while facilitating passive and targeted delivery. ^4^ Poly(lactic-co-glycolic acid) (PLGA) is a biodegradable and biocompatible polymer, that has emerged as a versatile platform for engineering nanoparticles for cancer therapy. ^5,6^ PLGA nanoparticles (NPs) encapsulating of PTX (PTX@PLGA), have demonstrated potential to enhance drug efficacy making them a promising strategy for cancer treatment. ^7–9^

Cellular metabolism plays a critical role in cancer progression and therapeutic response, where tumor cells exhibit metabolic reprogramming, characterized by the production of energy mainly by glycolysis, along with other factors that assist cancer cells to proliferate and survive. ^10,11^ From this point of view, evaluating metabolic changes in cancer cells in response to treatment can provide valuable insight into drug efficacy and acquired cellular resistance. ^12,13^ To assess cellular metabolism, multiphoton fluorescence lifetime imaging microscopy (MP-FLIM) has emerged as a powerful, label-free imaging tool.^13–16^ By exploiting the intrinsic autofluorescence of metabolic cofactors such as nicotinamide adenine dinucleotide (NAD(P)H), MP-FLIM provides quantitative information about the cellular redox state and balance between glycolysis and oxidative phosphorylation (OXPHOS), which are considered hallmarks in monitoring of cancer cells. ^16,17^ The MP-FLIM technique offers several advantages, including non-invasive monitoring with no need for external dyes, high spatial resolution, and the ability to measure dynamic metabolic changes in real time, crucial for continued monitorization.

In this study, we developed monodisperse PTX loaded PLGA (PTX@PLGA) NPs with sustained and controlled drug release profile. Their efficacy was tested and compared to that of free PTX and as a control the empty nanocarrier. Metabolic imaging based on label-free MP-FLIM of the metabolic cofactor NAD(P)H was used to assess the therapeutic response of monolayers of A549 lung cancer cells upon treatment with free PTX and PTX@PLGA NPs. Two established analysis methods were used for comparison on the obtained MP-FLIM data, namely the model-based curve fitting and model-free phasor plot approach. This work provides not only an understanding of the cellular metabolic response to nanoparticle-based drug delivery but also showcase label-free MP-FLIM as a valuable tool for assessing drug efficacy in cancer research.

## Materials and Methods

### Synthesis of paclitaxel-loaded nanoparticles PTX@PLGA NPs

PTX@PLGA NPs were prepared using the single emulsion solvent evaporation method. Briefly, an organic phase of 20 mg/mL of PLGA (39.5 kDa, Sigma-Aldrich) and 1 mg/mL of PTX (MedChemExpress) was added to an aqueous phase of 2% polyvinyl alcohol (PVA) (Sigma-Aldrich). The phases were ultrasonicated at 12 W for 2 min (Disintegrator Ultrasonic Mod.450, Branson) in an ice bath to form the emulsion. The emulsion was then immediately added to more PVA solution and left for 1.5 h under magnetic stirring to evaporate the organic solvent. To remove non-encapsulated drug and stabilizer, the nanoparticles were ultracentrifuged thrice at 30000 g for 20 min at 4ºC (OPTIMA XE-100, Isaza equipped with rotor TYPE 70 Ti, Beckman Coulter). In between centrifugation steps, the supernatant was discarded, and the pellet was resuspended in water at 4ºC. Finally, the suspended nanoparticles were freeze-dried (Lyoquest −55ºC Plus Eco, VWR) for 24 h. Preparation of non-loaded nanoparticles were done following the same protocol.

### Physicochemical characterization of NPs

#### Size, polydispersity index and zeta-potential

The hydrodynamic size, polydispersity index (PDI), and zeta-potential were measured using dynamic light scattering (DLS) (SZ-100Z Dynamic Light Scattering System, Horiba) with a sample concentration of 0.2 mg/mL in water at 25ºC.

#### Morphology characterization

The morphology of individual nanoparticles was evaluated by transmission electron microscopy (TEM) (JEM-2100-HT, JEOL, operating at 80 kV). A 10 µL sample size of nanoparticle suspension (0.02 mg/mL) was placed on 400 mesh grids (Ted Pella) and stained with 3% phosphotungstic acid (Electron Microscopy Sciences) for 30 sec. After staining, the sample was blotted to remove excess fluid and left to dry for 24 h.

#### Encapsulation efficiency and loading capacity

The encapsulation efficiency (EE) and loading capacity (LC) were obtained through high-performance liquid chromatography (HPLC). The HPLC system (Ultra-HPLC 1290 Infinity II LC System, Agilent) was equipped with a 2.6 µm, 150 mm × 4.6 mm C18 column (Kinetex), and PTX absorbance was measured at a wavelength of 232 nm.

Briefly, PTX@PLGA NPs was dissolved in 0.1% phosphoric acid (Sigma-Aldrich) in 50:50 acetonitrile (ACN, Sigma-Aldrich) and water mixture. The suspension was incubated for 6 h at 50ºC to allow polymer dissolution and release of PTX. The polymer precipitated by centrifugation at 20000 g for 30 min (Frontier 5718R, Ohaus). The supernatant with PTX was removed, filtered and analyzed.

The mobile phase consisted of water (solvent A) and ACN (solvent B). Linear gradient elusion was applied from 50% to 90% of solvent B in 9 min at 1.3 mL/min flow rate. A PTX calibration curve was pre-determined by injecting samples of known PTX concentration ranging from 0.2 to 450 µg/mL.

The EE and LC were calculated using Equations 1 and 2 respectively.

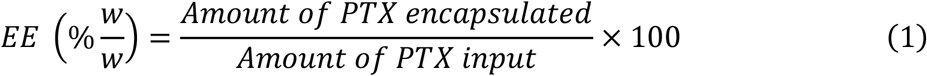

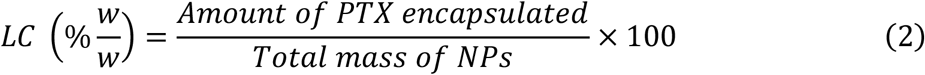

#### Drug release profile

The drug release profile of the PTX@PLGA was determined by dialysis diffusion to quantify the amount of PTX released throughout a 72-h time interval. 20 mg of NPs was dispersed in 2 mL of phosphate buffer solution (PBS, Corning) and accommodated in a dialysis membrane (14kDa, MediCell Membranes). The membrane was dialyzed against 40 mL of PBS at 37ºC, stirring at 300 rpm. At predetermined intervals of 0.5, 1, 1.5, 2, 3, 4, 5, 6, 8, 10, 12, 24, 36, 48, 60, and 72 h, 0.5 mL of the release media was collected, and 0.5 mL of ACN was added to the sample. The PTX released was quantified using the same HPLC protocol as previously described. The volume of media removed for the release quantification was replaced with new PBS after each sampling.

### *In vitro* cytotoxicity studies

#### Cell culture protocol

A549 cells were grown in Dulbecco’s modified eagle medium (DMEM, Biowest) supplemented with 10% fetal bovine serum (Hyclone) and 1% penicillin-streptomycin (PAN-Biotech) in 25 cm^2^ cell culture flasks at 37ºC in a humidified atmosphere with 5% CO_2_. Subcultivation was performed using 1:5 ratio and the media was renewed every 2-3 days, according to ATCC recommendations.

Spheroid cultivation was done in 96-well ultra-low attachment plates (faCellitate - BIOFLOAT™). 500 cells/well were seeded in phenol-red free DMEM supplemented with 10% FBS and 1% PS. The plate was then centrifuged at 1300 rcf for 3 min to accelerate the aggregation process of the cells. The cells were left in the incubator, and the spheroids were assessed on day 3 of cultivation. For metabolic imaging the spheroids were transferred to a µ-Slide 15 Well dish (Ibidi).

#### Determination of IC_50_

To determine the IC_50_, 10^4^ cells/well of A549 were seeded in 96-well plates. A range of concentrations of free PTX between 0.0005 - 5 µM, and 0.00015 - 1.5 mg/mL of PTX@PLGA NPs was tested. After 24 h, the treatment media was removed, and the cells were washed with PBS to remove any traces of the drug, and Resazurin (Sigma Aldrich) cell viability assay was carried out. Treatment media was removed from each well, and the cells were washed with PBS twice. Working solutions contained 10 µg/mL resazurin stock solution diluted in media, which was added to the monolayers and left incubated for 4 h at 37ºC. The fluorescence intensity was measured using a microplate reader (H1 Microtiter Plate Reader, Synergy Biotek) with a 560 nm excitation/590 nm emission filter setting.

#### Live/dead assay

Live/dead assay was performed using a laser scanning confocal microscope (Zeiss LSM-780). Treatment media was removed and replaced with staining media. The staining media consisted of a mixture of 2 µM of calcein-acetoxymethyl (Setareh Biotech, Ex/Em = 488/493-582 nm) and 4 µM of propidium iodide (Acros Organic, Ex/Em = 561 nm/582-718 nm) in phenol-red free DMEM (Gibco), and the monolayers were incubated for 10 min at 37ºC.

#### MP-FLIM instrumentation

For label-free multiphoton metabolic imaging a custom-built inverted microscope setup was used, as seen in figure S1. It comprises a femtosecond pulse laser excitation source with emission wavelength centered at 783 nm (FemtoFiber smart 780, Toptica, 85 fs pulse length, rep. rate 80 MHz). Pulse compression was performed using a single-prism (89-844, SF10, Edmund Optics) compressor configuration as described in Ref.: ^18^. After pulse compression, the laser enters a galvanometer scanning system (GVSK2-EC, Thorlabs) and beam movement is performed through a telecentric scan (SL50-CLS2, Thorlabs) and tube (TTL200-B, Thorlabs) lens system. The beam enters the two-layer inverted microscope (ECLIPSE Ti2, Nikon), where a silver mirror is mounted in the filter cube in the bottom layer, and a dichroic long pass filter (F47-639, AHF) is mounted in the filter cube in the second layer. The excitation laser is passing the long pass filter while the collected signal is reflected. The microscope is equipped with a motorized XY stage (MLS203-1, Thorlabs) for sample positioning and a Z-scanning stage (ND72Z2LAQ PIFOC, PI) for focusing. Imaging is performed with an air objective (40X / 0.75 NA/ WD 660 µm Plan Fluor, Nikon). The cells were kept in a top stage incubator at 37ºC at 5% CO2 during the experiments (TC-MWP, Bioscience Tools). In the detection path 3 short pass emission filters (F76-639, AHF) are mounted in series with a dichroic filter (F38-489, AHF) and an additional short pass filter (F37-492, AHF) to separate the NAD(P)H signal. The NAD(P)H fluorescence signal is focused on a PMT hybrid detector (HPM-100-40; Becker & Hickl) which is connected to a TCSPC FLIM 150N module (Becker & Hickl GmbH), allowing characterization of the NAD(P)H fluorescence lifetime. The pulse synchronization output of the laser was used for time synchronization between laser pulse and the TCSPC card. Data was acquired through the SPCM software version 9.86 (Becker & Hickl GmbH, Berlin, Germany).

Fluorescence decay histograms were acquired with 1024-time bins in a time window of 12.5 ns and with a temporal resolution of 0.01 ns. A pixel dwell time of 5 ms was used for images of 256 × 256-pixels covering a 100 × 100 µm scan area, with a scan pixel resolution of 0.39 µm.

### Data analysis

#### Curve fitting

The FLIM data was processed using SPCImage software version 8.9 (Becker & Hickl GmbH, Berlin, Germany). A spatial square binning of 2 was applied to the images to enhance the signal-to-noise ratio and improve the fitting quality, while the curves were fitted using the maximum likelihood estimation (MLE) fitting algorithm. Considering only the fluorescence lifetimes of free and bound state of NAD(P)H, the data was fitted to a double exponential function () as seen in Equation 3.

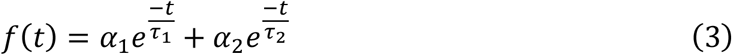

Where τ_1_ and α_1_ correspond to the fluorescence lifetime and preexponential factor (amplitude), respectively, of free NAD(P)H and τ_2_ and α_2_ correspond to the fluorescence lifetime and preexponential factor, respectively, of bound NAD(P)H. From the weighted contributions of the two fluorescence lifetimes a mean fluorescence lifetime τ_m_ for NAD(P)H is calculated. τ_m_ is a simple metric to calculate and is sensitive to changes in the environment but may obscure specific metabolic shifts by blending free and bound populations, we therefore also calculate a second quantity, namely the metabolic index (MI). MI is calculated as the ratio between the preexponential factors for the free and bound NAD(P)H (Equation 4) and was used to evaluate the cell’s redox state by quantifying the balance between metabolic pathways (e.g., glycolysis vs. OXPHOS).^19,20^

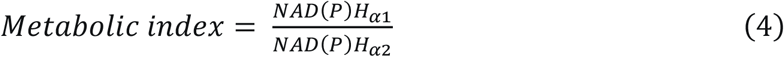

The acquired data are presented as colormaps for the NAD(P)H τ_m_ and MI, respectively.

#### Phasor analysis

To represent the obtained fluorescence lifetime data in a model-free approach a custom Python script was developed to extract lifetime components and phasor-derived metrics from the FLIM measurements, following the method described in Ref:.^21^ This approach apply total least square analysis on the phasor plane together with a 95% confidence ellipse to estimate lifetime parameters without requiring multi-exponential curve fitting. The software script also includes configurable pre-processing inputs (e.g., laser repetition rate, thresholds) and steps such as noise removal and temporal offset correction. The current phasor plot implementation does not allow for correlation between image pixel and data point in the phasor cloud, this feature could be implemented in future versions but is beyond the scope of this study.

#### Statistical analysis

Statistical analysis was performed using Origin software version 9.00 (OriginLab Corporation, Northampton, MA, USA), which uses one-way analyses of variance (ANOVA) or two-way ANOVA. Statistically significant differences were considered for *p<0.05, **p<0.01, ***p<0.001, ****p<0.0001. Results are shown as arithmetic means ± standard deviation.

## Results and discussion

### Physicochemical Characterization of PTX@PLGA NPs

The PTX@PLGA NPs were prepared by emulsion-solvent evaporation, chosen based on PTX’s drug solubility. Since PTX is a hydrophobic compound, it can be entrapped in the hydrophobic core of PLGA NPs.

The hydrodynamic size, polydispersity index (PDI), and zeta potential values of the empty PLGA and PTX@PLGA NPs, measured using DLS, are represented in Table 1.

**Table 1:** Physicochemical characteristics of PLGA and PTX@PLGA NPs.

|  | *Hydrodynamic*<br>*size (nm)* | *Polydispersity*<br>*index* | *Zeta-*<br>*potential*<br>*(mV)* | *Encapsulation*<br>*efficiency (%)* | *Loading*<br>*capacity*<br>*(%)* |
| --- | --- | --- | --- | --- | --- |
| PLGA | 202.91 ± 4.50 | 0.10 ± 0.03 | -30.07 ± 6.88 | n.a. | n.a. |
| PTX@PLGA | 219.43 ± 9.17 | 0.11 ± 0.06 | -41.73 ± 9.59 | 30.50 ± 5.40 | 1.39 ± 0.60 |

n.a: not applicable

Compared to empty NPs, PTX@PLGA NPs showed a larger hydrodynamic size, suggesting that the loading of the nanoparticles was successful. Moreover, the obtained size of the NPs diameter was considered appropriate for the enhanced permeability and retention effect within tumor microvasculature. ^22^ The PDI of both PLGA and PTX@PLGA NPs was lower than 0.2, indicating a narrow size distribution and a monodisperse population. ^23^ Moreover, the PDI did not increase with the drug loading, suggesting that the drug encapsulation method was successful in the formulation of a monodisperse population of drug-loaded NPs. ^8^ The obtained zeta potential values below −30 mV for both types of NPs indicate that the nanoparticles achieved colloidal stability by electrostatic repulsive forces that prevent nanoparticle aggregation. ^24^, where the negative value can be attributed to the terminal carboxylate group of the PLGA polymer. ^25 24^

The morphology of PTX@PLGA NPs after the freeze-drying process was observed via TEM, as seen in figure 1a. The NPs presented a spherical morphology of around 68 nm, where the spherical shape is reported to be more biocompatible than other types of morphologies. ^26^

**Figure 1:**
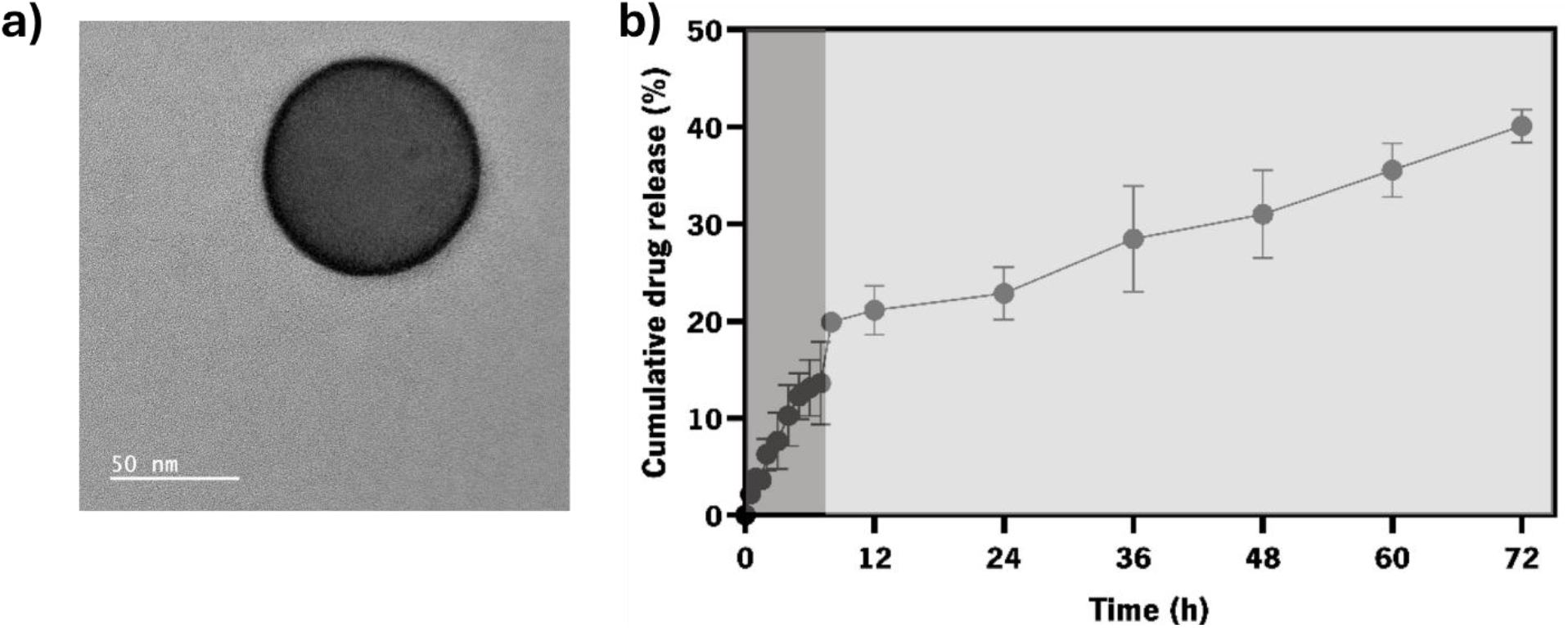
Physicochemical characterization of PTX@PLGA NPs. a) Representative TEM image of a single PTX@PLGA NP showing an average diameter of 68 nm. b) Cumulative amount of PTX release from the PTX@PLGA NPs between 0 and 72 hours.

The last two columns in Table 1 summarizes the encapsulation efficiency (EE) and loading capacity (LC) of PTX@PLGA NPs calculated by equations 1 and 2. The EE indicates that 30.5% of the PTX initially used in the formulation process was successfully encapsulated within the PLGA NPs. This level of encapsulation suggests moderate efficiency. Meanwhile, the LC reflects the amount of PTX loaded relative to the total weight of the nanoparticles, indicating that 1.39% of the NP mass is comprised of the drug. This relatively low LC implies that while the nanoparticles encapsulate the drug, the proportion of drug to carrier is small. Nonetheless, the EE and LC agree with previous studies using the synthesis protocol mentioned. ^25^

The PTX@PLGA NPs showed a two-phase release pattern, as seen in figure 1b, characterized by an initial burst over the first 8 h and a sustained release up to 72 h.

The initial rapid release of PTX can be attributed to the diffusion of the drug that is adsorbed to the nanoparticle surface and was not encapsulated, or the PTX near the surface due to drug redistribution after the freeze-drying process. ^27^ Nonetheless, burst release is beneficial for achieving therapeutic concentration in minimal time. ^28^ The second phase of release can be explained by the degradation of the PLGA matrix and subsequent diffusion of PTX from the core. The sustained release ensures a prolonged therapeutic effect, reducing the need for frequent dosing and potentially minimizing the side effects associated with high systemic drug concentrations, such as nausea, neutropenia, anemia, neuropathy, hair loss, and potential long-term damage to other organs, and by such this two-phase pattern is advantageous, allowing for both immediate and long-term action. ^29^

### Cytotoxicity of free PTX and PTX@PLGA nanoparticles

The cytotoxic effects of PTX and PTX@PLGA NPs on A549 monolayers were evaluated to determine their respective IC_50_ values. First, empty PLGA NPs were found to be non-cytotoxic for concentrations up to 5mg/mL, as seen in figure 2a. No significant statistical difference compared to the control, and no values below 70% threshold, which is in accordance with the International Organization for Standardization (ISO) 10993-5.

**Figure 2:**
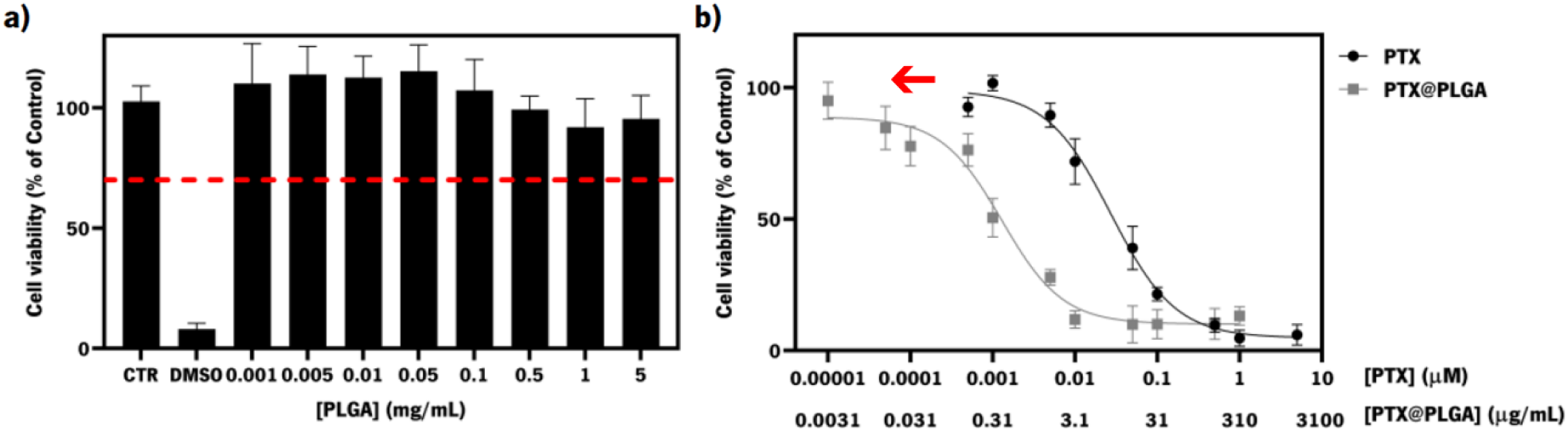
Cell viability assay and IC_50_ determination. a) Cell viability for different concentrations of empty PLGA nanoparticles compared to the control. b) Dose-response curve of free PTX and PTX@PLGA nanoparticles on monolayers of A549 cells.

The dose-response curve for PTX and PTX@PLGA NPs, as represented in figure 2b, shows a decrease in cell viability with increasing drug concentrations, indicating a typical sigmoidal relationship between drug concentration and cytotoxicity. From the dose-response curves presented, the IC_50_ after 24 hours of incubation for free PTX and PTX@PLGA NPs was determined to be 0.028 µM and 0.94 µg/mL, respectively. The calculated IC_50_ concentration of PTX, when encapsulated, was found to be 0.0013 µM, which is more than 20 times lower and considered significantly lower (p<0.0001) when compared to the IC_50_ concentration of free PTX. The effect is clearly observed by the notable leftward shift of the dose-response curve of PTX@PLGA, compared to the curve for the free PTX. These findings indicate that encapsulation of PTX enhanced anticancer activity compared to the free drug, as observed before in other studies. ^30,31^

The cytotoxicity of PTX, empty PLGA, and PTX@PLGA NPs was further evaluated by live/dead assay immediately after adding the formulation (0 h) and after 6 h, 12 h, and 24 h of administration, as shown in figure 3. The A549 monolayers were treated with the determined IC_50_ concentration of PTX and PTX@PLGA NPs. The different timepoints correlate roughly with the middle point of the burst release (6 h), the end of the burst release and the start of the slow release (12 h), and during the slow release (24 h). The treated PTX and PTX@PLGA NPs samples were compared to two different controls of untreated and empty PLGA NPs. To ease the interpretation of the outcome of each imaging, pie-charts inserts indicating the live (green) and dead (red) cells of the evaluated pixel in each image were added.

**Figure 3:**
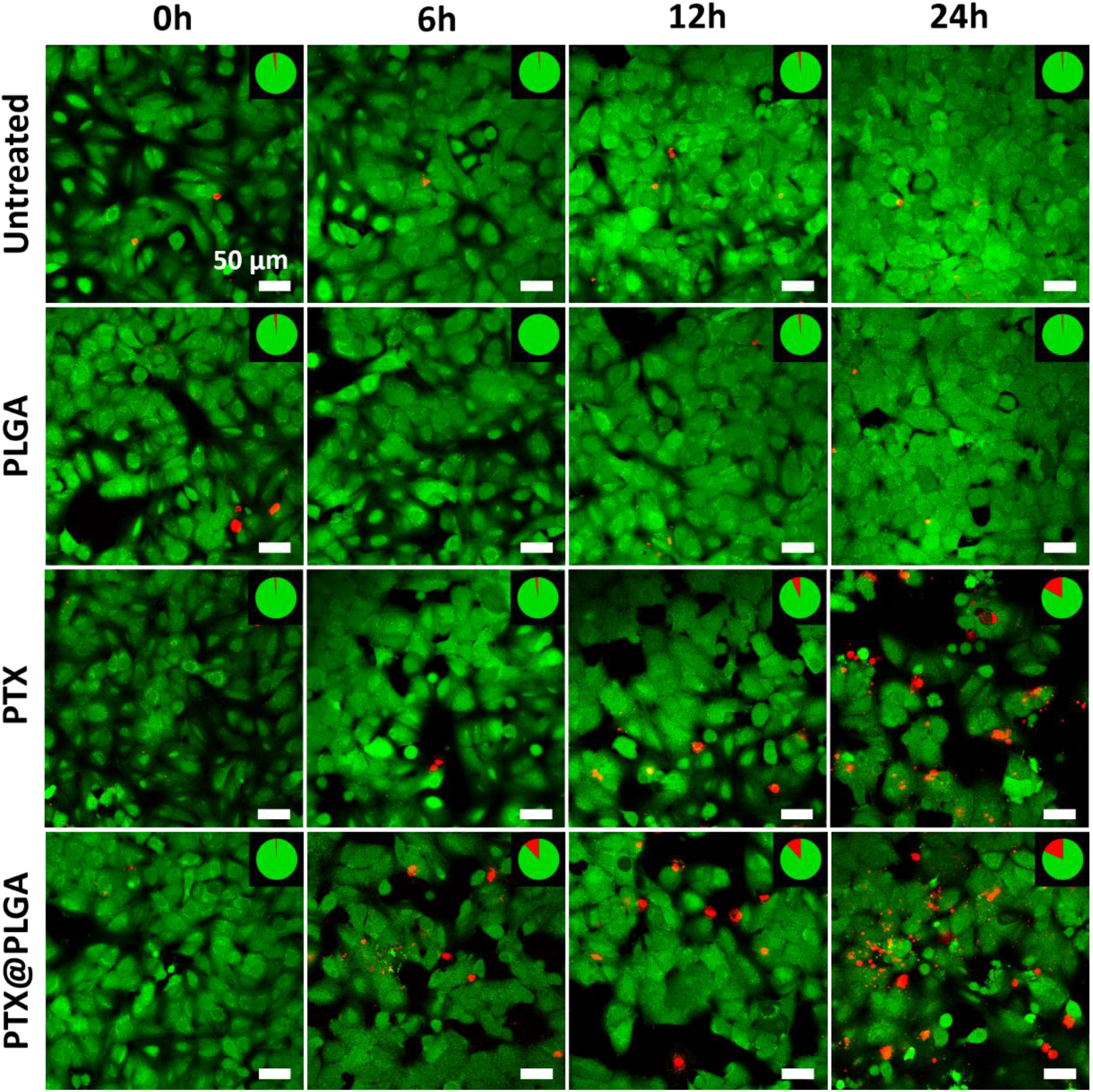
Live/dead assay of treated and non-treated A549 cells. Confocal fluorescence images taken on A549 cells showing untreated, and treated with empty PLGA NPs, PTX and PTX@PLGA NPs (from top to bottom), at the time points of 0, 6, 12, and 24 h (from left to right) after administration. Insets show the ratio of dead (red) to live (green) in a pie chart. Scale bar 50 µm.

For all samples at 0 h, the A549 monolayers exhibited a predominantly viable cell population, as indicated by the green fluorescence staining. The live/dead ratio was close to 100%, confirming the cells’ healthy status before treatment in all conditions, further indicated by the full green pie chart inserts in each subfigure at 0 h. Untreated cells maintained a high live/dead ratio, and cell confluence increased by 24 h following the approximately 22 h doubling time of A549 cells. ^32^

The empty PLGA-treated group showed no difference compared to the untreated cells, with a live/dead ratio of almost 100%, which is consistent with the previous knowledge presented in figure 3a that there is no cytotoxic effect at this concentration of empty NPs. Treatment with PTX showed an apparent increase in non-viable cells at 24 h, with a small or no considerable appearance of dead cells prior to that. The PTX@PLGA-treated group showed a moderate increase in dead cells at 6 h compared to the controls and PTX-treated cells, suggesting an earlier cytotoxic response due to rapid cellular uptake of the NPs in combination with the initial burst observed in the release profile in figure 1b. By 24 h, the number of non-viable cells was comparable to the results found for the PTX-treated cell sample in accordance with the IC_50_ concentrations used.

### Metabolic response of A549 cells to treatment

To assess the therapeutic response of both free and encapsulated PTX a time-lapse study was performed involving the same time points as selected in the live/dead assay. The metabolic response of A549 monolayers to PTX, empty PLGA, and PTX@PLGA NPs, as well as the control (untreated) was assessed by label-free MP-FLIM of the metabolic cofactor NAD(P)H. Both curve fitting and phasor analysis approaches were used in the analysis. For comparison between the two analysis methods the τ_m_ (Eq. 3), and the MI (Eq. 4) are determined at each time point for each condition. Before any calculations a threshold was applied to each image pixel to eliminate background pixels. For each of the 4 conditions 3 different samples were imaged at 3 random locations on each sample were followed over 24 h. Representative images depicting τ_m_ is shown in figure 4a, as well as the MI for the same images in figure 4b for the curve fitting method which allows for the identification of individual cellular response. For the selected images in figure 4a and b phasor clouds are generated, as seen in figure 4c. The statistical analysis of all collected data is represented for the curve fitting method and phasor approach in figure 4d and e, respectively.

**Figure 4:**
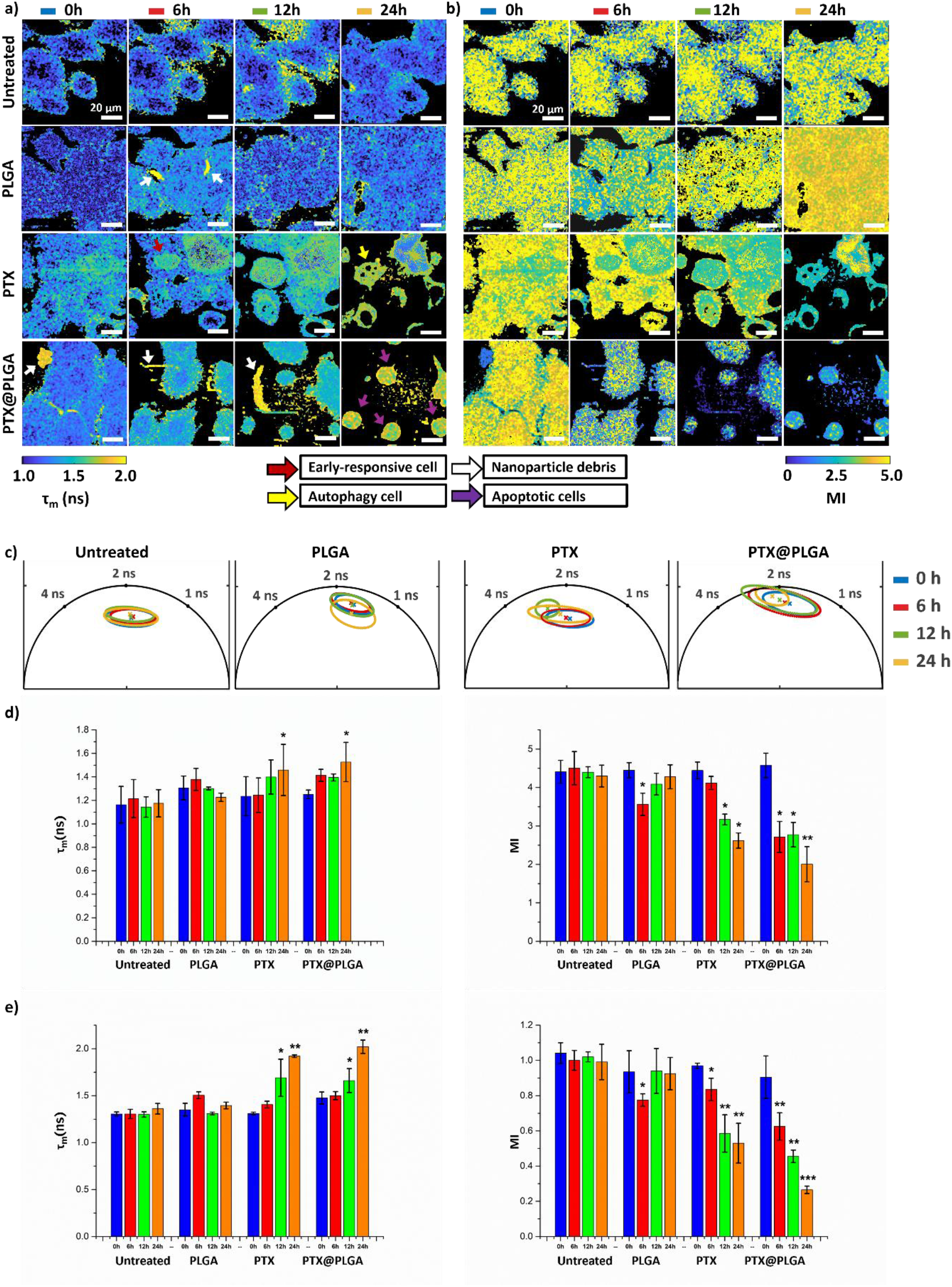
Label-free MP-FLIM therapeutic study of A549 cells including controls analyzed by both curve fitting method and phasor plot approach. a) Mean fluorescence lifetime (τ_m_), and b) Metabolic index (MI) both calculated using the curve fitting method for selected images. c) Phasor plots showing the extracted phasor clouds fit data for the different time points. d) Statistical analysis of τ_m_ and MI for the curve fitting method. e) Statistical analysis of τ_m_ and MI for the phasor approach. For d) and e) ^∗^p < 0.05, ^∗∗^p < 0.01, and ^∗∗∗^p < 0.001 compared to 0 h. Scale bar in a) and b) 20 µm.

For the untreated cells the image data in the first row of figure 4a and b shows small variations of the τ_m_ and MI values for different intercellular locations across different time points but does not correlate to a statistically significant variation as seen in analysis in figure 4d. In the statistical analysis of the MI in figure 4d, all conditions presented high values (>4) of MI, associated with the prevalence of glycolysis, which can be explained by the Warburg effect and the preference of glycolysis as main pathway for energy production. ^10^ These results are collaborated by the findings from the phasor method where the center (τ_m_) and shape (95% of the data points) of the phasor cloud seen in figure 4c stays constant across the time points and the extract value for τ_m_ is comparable to the one found by the curve fitting method. The absolute value of the extracted MI for the two methods differs, which is not significant for the result of the analysis and both methods show the same trends and behavior for the control.

For the second type of control sample of empty PLGA-treated cells the images of τ_m_ and especially the MI show, in the second row of figure 4a and b differences after 6 h of incubation but no morphological changes of the cells are observed, suggesting a metabolic response to the nanocarrier itself. This may be due to the degradation of the PLGA matrix into lactic and glycolic acid which are metabolized through the Krebs Cycle. ^33^ The PLGA-treated cells displayed a 19% significant decrease (p<0.05) in the MI at 6 h, followed by an increase back to the initial value, indicating the cells recover the initial more glycolysis-based metabolism. The phasor clouds in figure 4c for the empty PLGA-treated cells do not indicate a change at 6 h and only a pure vertical translation of the phasor cloud is observed at 24 h, which could be indicative of a change in the background signal. However, the change is reflected in the statistical analysis for both methods where a slight increase in the τ_m_ is observed after 6 h, decreasing again by 12 and 24 h. The MI shows the same trend with a significant decrease after 6 h and the return to base value at 12 and 24 h. The translation of the phasor cloud at 24 h does not seem to affect the values of τ_m_ and MI as seen in figure 4e.

Additional for the empty PLGA and PTX@PLGA-treated cells conditions, small features (marked with white arrows in figure 4a) with a significant higher τ_m_ value can be observed and are associated with debris from the NPs synthesis process. The associated pixels were identifying and removed in the curve fitting analysis as seen in figure S2. In the phasor approach, there is no possibility to identify and correlate the individual pixel position, and we cannot remove the pixels related to the NPs debris, for the statistical analysis of τ_m_ and MI. In future iterations of the custom software this functionality could be added but is beyond the scope of this work. However, no observable difference is seen in the statistical analysis between the curve fitting method and the phasor approach.

In the third row of figure 4a and b PTX-treated cells presented metabolic heterogeneity at 0 h. After 6 h of treatment, different τ_m_ and MI values are observed across the sample, indicating changes in the metabolic state of the cells. An increase in τ_m_ indicates a shift towards OXPHOS and can be observed by the localized cyan colored pixels suggesting an early response to the drug, marked by a red arrow. After 12 h, an increase in τ_m_ across all cells is observed, as well as the rounding of cells, indicating cells are dying and detaching from the plate. After 24 h, the τ_m_ continued to increase, indicated by the prominent green and yellow color. Besides the change in τ_m_, the early-responsive cell showed significant morphological changes. The appearance of vacuolization is observed after 24 h, marked by yellow arrows which can be associated with autophagy, a self-renewal process where the cell cleans out damaged proteins and cellular structures and being considered as a survival mechanism. ^34^ Autophagy has been previously reported in A549 cells in response to PTX and linked to drug resistance. ^35–37^

In figure 4d the PTX-treated monolayers suffered a gradual decrease over time in the MI value, showing a small but not significant (p>0.05) lowering at 6 h, a significantly lowering (p<0.05) at 12 h, and by 24 h it further decreased to 41% of its initial value (p<0.05). The decrease in MI indicates that the contribution of bound NAD(P)H increased, and the cells metabolically shifted towards OXPHOS as response to the administration of PTX. The phasor cloud for the PTX-treated cells in figure 4c shows no change at 6 h, and a horizontal leftwards translation of τ_m_ towards longer lifetimes at 12 h, indicating a shift towards OXPHOS. After 24 h the phasor cloud showed an elongated shape as compared to the initial conditions indicating that the cells are changing metabolic states. The statistical analysis for the phasor approach shows the same overall tends as seen for the curve fitting method but seems to be more sensitive showing a larger and more significant shift for both τ_m_ (24 h) and MI (12 and 24 h).

For the PTX@PLGA-treated cell condition, fourth row of figure 4a, an increase in the τ_m_, accompanied by morphological changes are observed. By 12 h, cells had already fully detached from the plate, indicating cell death, and their metabolism continuously shifted as apoptosis was occurring. ^38^ By 24 h, the cells show clear signs of apoptosis with formation of apoptotic bodies, marked by purple arrows with high τ_m_ values. The PTX@PLGA-treated cells displayed, as for the PTX samples a shift of τ_m_ to longer lifetimes associated with a shift towards OXPHOS. The MI calculated for the PTX@PLGA-treated cells displayed a significant decrease already after 6 h (p<0.05), due to the fast internalization of the PLGA NPs, as corroborated with the results found in the live/dead assays (figure 3). The MI was observed to maintain a constant value from 6 to 12 h (p<0.05) and by 24 h had decreased further to 55% of its initial value (p<0.01). The difference in the decrease between free and encapsulated PTX can be due to the drug release profile of the NPs. The initial decrease could be associated with the burst release of the PTX that happens in the first 8 h, while the controlled and sustained release leads to more subtle changes in the MI. The phasor cloud displays a translation of τ_m_ to longer lifetime and elongation of the ellipse corroborating the findings from the curve fitting method. Comparing the extracted values for τ_m_ and MI again showing larger relative changes, suggesting that the phasor approach was more sensitive in detecting more subtle metabolic changes.

Overall, A549 lung cancer cells shifted to OXPHOS in response to PTX-induced cell death as expected since PTX induces apoptosis by interfering with the cell cycle, as reported previously in this cell line. ^39–41^ Furthermore, other metabolic imaging studies on different cancer types, PTX has been shown to induces a shift to OXPHOS ^42,43^. Furthermore, the response to a drug delivery system on a single A549 cell level has been previously evaluated using MP-FLIM, however, the influence of the controlled release by the nanoparticles and the metabolic response to the carrier itself was not studied. ^44^

In summary, curve fitting method provides precise quantitative analysis, however, it can be affected by the model and initial guesses and is computationally intensive. Phasor plots proved to be slightly more sensitive to record metabolic changes in our study and provide a rapid screening and visual representation of the raw FLIM data. ^45^ Overall, both data analysis methods reveal a shift in the metabolic activity to OXPHOS in response to the chemotherapeutic drug PTX. PTX@PLGA NPs displayed a higher metabolic shift in both analysis methods as compared to the free PTX, indicating that this type of treatment modality has great benefits comparing to the free drugs.

### Metabolic imaging in 3D

In the last section we show preliminary results extending the extraction of metabolic data from 2D cellular monolayers to a 3D spheroid model. For the experiment a mono-culture spheroid of A549 cells with a diameter of 270 μm was imaged after 3 days of cultivation. These kinds of compact spheroids are known to have nutrient and oxygen gradients that lead to the sectioning of spheroids into proliferating cells, quiescent cells, and a necrotic core, consequently leading to different metabolic states. ^46–48^ The concept of the experiment was that the spheroid would show signs of metabolic differences as a function of distance from the center. An image stack was therefore created from the spheroid’s tip touching the plate’s bottom (z = 0 μm), to roughly the center (z = 140 μm), as represented with the blue square in figure 5a. Three selected image depths of 0, 70 and 140 μm corresponding to the outer layer, halfway to center and the center, respectively are shown for τ_m_ and MI in figure 5b. The full image stack with a step size of 10 μm for τ_m_ and MI are shown in figure S3a and b, respectively. The colormaps of both τ_m_ and MI in figure 5b and figure S3a and b, were obtained by the curve fitting method.

**Figure 5:**
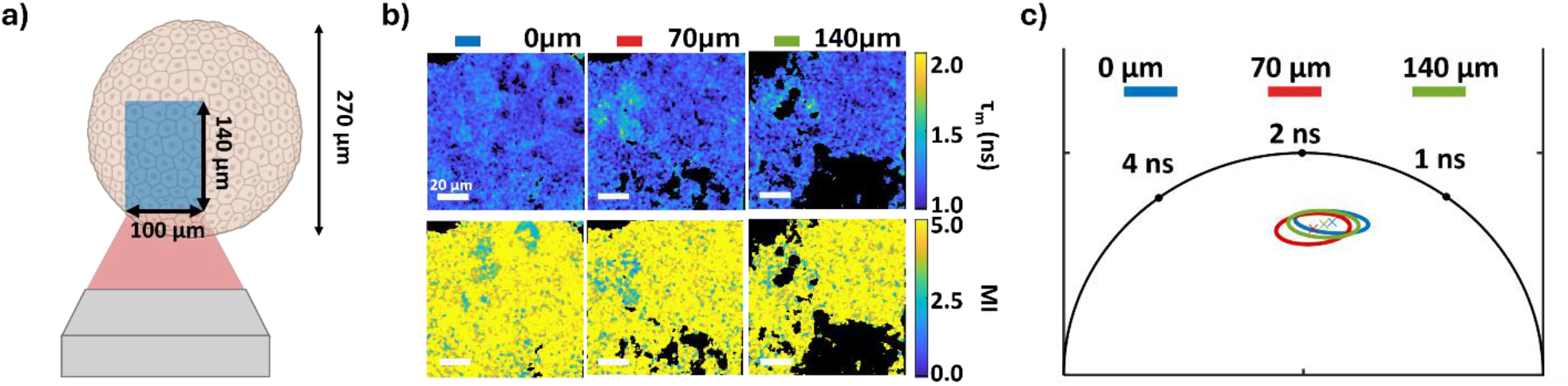
3D metabolic imaging. a) Schematic representation of the A549 spheroid and selected 3D imaging stack. b) Multi-photon FLIM images represented by τ_m_ with a z step size of 10 µm (Scale bar 20 µm). c) Phasor representation of τ_m_ for 3 selected image heights (0, 70, and 140 µm).

The overall value for τ_m_ in each image plane of the stack is similar the values found for the control condition for the monolayers, however there are areas indicating a higher τ_m_ values which are observed close to dark areas which are correlated to no metabolic activity (no fluorescence signal) or voids in the spheroid. A similar behavior is found in the MI colormaps showing mostly high MI values, as found in the monolayer controls. The areas indicated to have a higher τ_m_ are even more evident in the MI images showing lowered values equivalent to the observations in monolayers for the conditions treated with PTX and PTX@PLGA NPs and associated with OXPHOS. The large dark area observed in the lower right corner for deeper image depths in figures 5b and S3 could be linked to a necrotic core in the spheroid.

The phasor plot analysis corroborates these findings, figure 5c represents the phasors correspondent to the selected tissue depth at 0, 70, and 140 μm. No shifts in the phasor cloud are observed comparing the outer layer (0 μm) with the deepest layer (140 μm), suggesting no metabolic distinction between the edge and centre of the spheroid. A slight leftwards shift at 70 μm is observed, however it may be due to the presence of a larger number of cells that show a lower value of the MI as discussed above.

Overall, small metabolic differences were observed, suggesting that the spheroids of imaged size do not fully mimic the tumor microenvironment, and that spheroid development must be optimized to include more tumor microenvironment features, including more pronounced metabolic gradient. For that reason, studies with different cell densities and longer cultivation days are needed to obtain compact and metabolically differentiated spheroids.

## Conclusion

This study highlights the utility of label-free MP-FLIM for evaluating the metabolic responses of A549 lung cancer cells to PTX and PTX@PLGA NPs. Our findings demonstrate that PTX@PLGA NPs induce a pronounced metabolic shift toward OXPHOS, with a greater degree of change compared to free PTX. This enhanced metabolic response correlates with the sustained and controlled drug release profile of the NP formulation, underscoring the advantages of nanocarrier-based drug delivery systems in achieving therapeutic effects at low dose concentrations.

Metabolic imaging proved to be promising in the study of therapeutic response on the cellular and subcellular level by tracing treatment-induced metabolic shifts as the cancer cells adapt metabolically to the therapy. In our analysis of the metabolic shifts, we employ both the model-based curve fitting method and the model-free phasor plot approach, where both methods reach comparable results in the case of monitoring the single metabolite NAD(P)H and its two forms. We further show that by employing label-free MP-FLIM metabolic data from 3D models, are possible, strengthening the general applicability of the method to more advanced models, including patient-derived models.

In summary the results not only emphasize the potential of PTX@PLGA NPs as an improved treatment strategy for lung cancer but also validate MP-FLIM as a powerful tool for investigating drug-induced metabolic dynamics. These insights pave the way for further exploration of NP formulations and their role in modulating cellular metabolism, offering new perspectives for the optimization of cancer therapeutics.

## Supporting information

Complete supplemental material and TOC figure

## Conflict of interests

The Authors declare no conflict of interest.

## AUTHOR INFORMATION

### Present Addresses

†If an author’s address is different than the one given in the affiliation line, this information may be included here.

### Author Contributions

The manuscript was written through contributions of all authors. All authors have given approval to the final version of the manuscript.

Lia Santos:

Conceptualization (equal), Data curation (lead), Formal analysis (lead), Investigation (equal), Methodology (equal), Validation (equal), Visualization (equal), Writing – original draft (lead), Writing –review & editing (equal)

Maria Leonor Ribeiro

Conceptualization (supporting), Data curation (equal), Formal analysis (equal), Investigation (supporting), Methodology (equal), Writing – review & editing (supporting)

Jana B. Nieder:

Conceptualization (equal), Formal analysis (supporting), Funding acquisition (equal), Investigation (equal), Methodology (equal), Project administration (equal), Resources (equal), Software (equal), Supervision (equal), Validation (equal), Visualization (supporting), Writing – original draft (supporting), Writing – review & editing (equal)

Christian Maibohm:

Conceptualization (lead), Data curation (equal), Formal analysis (equal), Investigation (equal), Methodology (equal), Validation (equal), Visualization (equal), Writing – original draft (lead), Writing –review & editing (equal)

### Funding Sources

We thank the funding agencies CCDR-N and FCT for funding via the UTAustin-Portugal program, for the project ‘ExtreMed’ with grant no: NORTE-01-0247-FEDER-045932 andHfPT – Health from Portugal, with the reference n.ºC644937233-00000047, co-funded by Component C5 – Capitalisation and Business Innovation under the Portuguese Resilience and Recovery Plan, through the NextGenerationEU Fund.

### Notes

Data statement:

Custom Python code for the phasor analysis is available upon reasonable request.

## ACKNOWLEDGMENT

The authors would like to thank Pedro Silva for the TEM imaging.

Maria Leonor Ribeiro would like to acknowledge *Fundação para a Ciência e a Tecnologia* (FCT) for the financial support through PhD grant 2022.13371.BD.

We thank HfPT Health from Portugal with reference number: n.ºC644937233-00000047 and the ExtreMed’ project with grant no: NORTE-01-0247-FEDER-045932for the support for this study.

## Notes

### Competing Interest Statement

The authors have declared no competing interest.

