## Supplementary material for "*In Vitro* Efficacy of Paclitaxel-loaded PLGA Nanoformulations for Lung Cancer Treatment Demonstrated by Label-free Multiphoton-Fluorescence Lifetime Imaging Microscopy": Complete supplemental material and TOC figure

Figure S1:

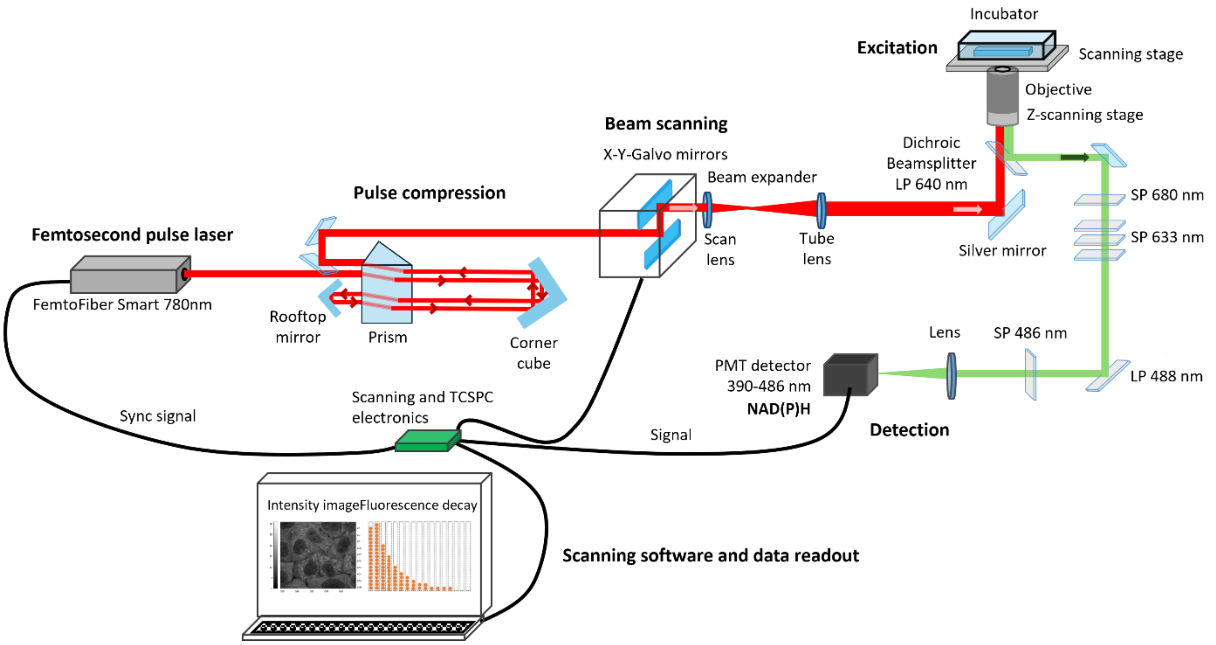


*Figure S1: Schematic representation of custom-built multi-photon microscope platform for metabolic imaging of NAD(P)H.*

Figure S2:

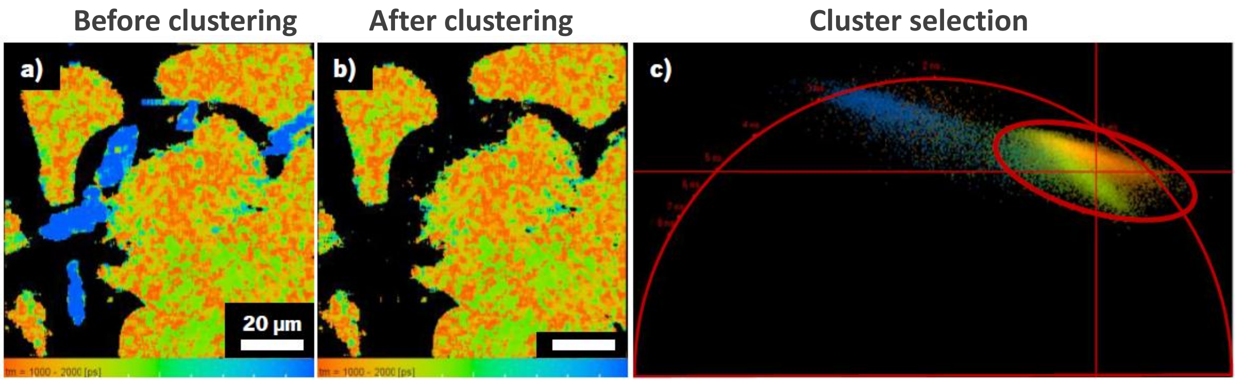


*Figure S2: Clustering was performed using phasor plots generated by the data analysis software to eliminate the fluorescence contribution from artifacts associated to debris from the lyophilized NPs (Figure S2). The cluster corresponding to the cells was selected, and the values of $\tau_m$, $\alpha_1$, and $\alpha_2$ were retrieved for statistical analysis (Figure S2).*

Figure S3;

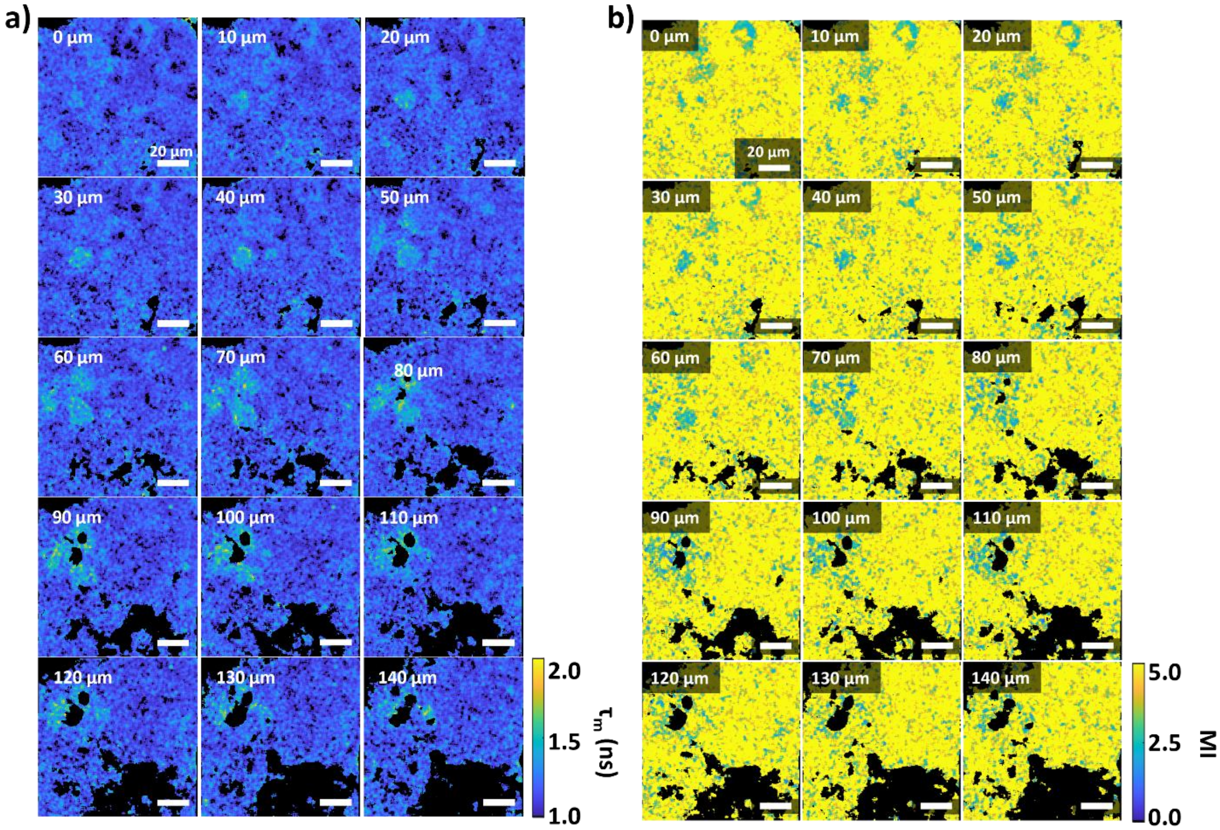


*Figure S3: Full image stack of 3D spheroid from 0 µm to 140 µm in steps of 10 µm. a) Mean fluorescence lifetime ($\tau_m$) of 3D spheroid as a function of image depth. b) Metabolic index (MI) of 3D spheroid as a function of image depth. Scale bar 20 µm.*

TOC figure:

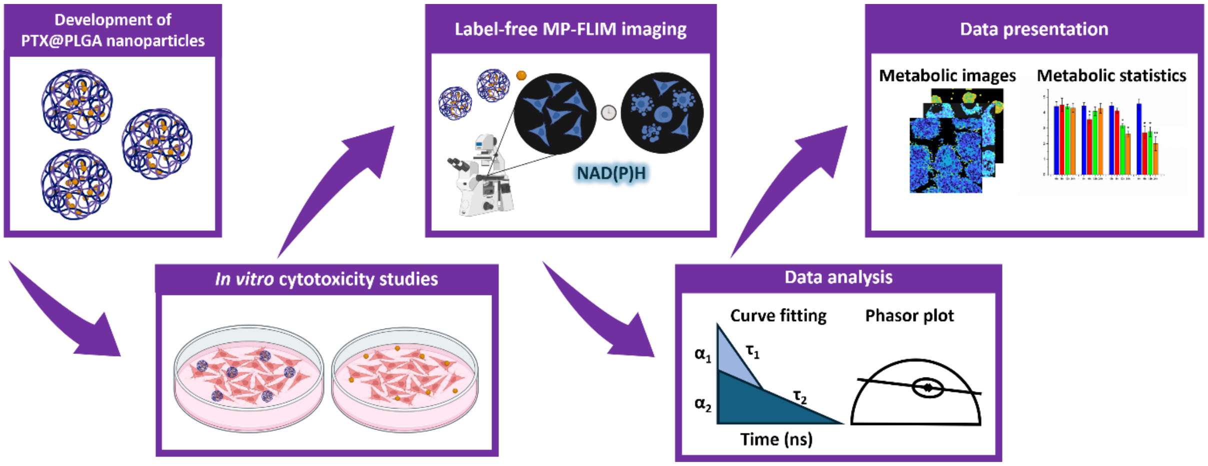


*TOC: Graphical representation of the presented study, including the development of the PTX encapsulated PLGA nanocarriers, in vitro studies, and metabolic multiphoton FLIM imaging with subsequent analysis through curve fitting and phasor plot methods. (created in Biorender.com)*
